# The Identification of A ROS Responsive Motif that is regulated by snoRNP in *Arabidopsis*

**DOI:** 10.1101/2020.06.08.141143

**Authors:** Han Cheng, Mingan Sun, Dianjing Guo

## Abstract

A GGGCC motif (site II like motif) was identified from the upstream sequences of reactive oxygen species (ROS) and light induced genes in Arabidopsis. This motif is highly enriched within the −50 to −250 bp region of the induced genes, and it is also specifically distributed in the same region in mouse and human genome. EMSA experiments revealed that several nuclear factors (NFs) bind to this motif, and the binding activities altered under H_2_O_2_ treatment. Two C/D family snoRNP proteins, namely fibrillarin 2 and NOP56, were identified from the site II like motif binding NFs. Several C/D family snoRNA, including R63, U24a and Z15, were also cloned from the motif binding NFs. These data suggest new regulatory roles of snoRNP in Arabidopsis.

## 1 Introduction

More than twenty percent of the earth’s atmosphere is composed of oxygen, which benefits most organisms on the planet as the final receptor of electron transport during respiration [1]. The oxygen is then turned into reactive oxygen species (ROS), e.g. superoxide (O_2_^·-^). ROS are quite unstable and prone to oxidize the biological molecules nearby, which may cause damages to the cell. There are several forms of ROS, which are quite dynamic and can interconvert quickly [2,3]. For example, superoxide is usually rapidly dismutated to hydrogen peroxide (H_2_O_2_), which is relatively stable and able to get across membrane [4,5]. Protonation of O_2_^·-^ yields hydroperoxyl (HO_2_^·-^) radical, which can attach to fatty acid directly [6]. In the presence of transition metal ions, such as Fe2+, H_2_O_2_will be turned into the most destructive hydroxyl (OH^·-^) radical [7]. In plant cells, there are two major sources for ROS production: leakage from electron transfer during respiration and photosynthesis [8]; or synthesis by a variety of enzymes, including peroxidases and oxidases [9–12]. The produced ROS are either degraded by catalases (CAT) [13–15] and sorts of peroxidases [16,17], or nicely controlled in particular spaces such as peroxisomes [18,19] to avoid possible damages to the membrane system and biological macromolecules. Antioxidants, such as ascorbic acid, tocopherol, uric acid and glutathione also play important roles in protecting cells from the damage of ROS [20,21].

Apart from the deleterious effects [22], ROS are also known as signal molecule involving in gene expression regulation [12,20,23]. Previous reports suggested that H_2_O_2_ are involved in ABA, GA, auxin and other hormone signal pathway [24–27], and thus participate in the regulation of particular physiological process and developmental program [25,28–31]. *Arabidopsis* Microarray data analysis revealed that about 1-2% of the transcriptome [32] and one third of the transcription factors were altered in mRNA expression after exposure to H_2_O_2_ [33]. In *Arabidopsis*, hundreds of genes are annotated as “ROS signalling related” [34–36]. Recently, an LRR receptor kinase HPCA1 was identified as H_2_O_2_ in Arabidopsis [37], however, very few ROS responsive DNA motifs were experimentally verified in plants [38–41]. Besides, the transcription factors that regulate the motif are yet to be identified. Because ROS have been proved to be universal signal molecules in plants, animals and other kingdoms, it is not clear whether a conserved regulatory mechanism exists across kingdoms. Identification of more ROS responsive motifs and the corresponding regulators, especially those potentially conserved ones, is therefore critical for elucidating ROS mediated signalling pathway.

Compared with animal cells, plant cells are more robust to ROS and contain relatively higher level of H_2_O_2_ [8,42,43]. In the green plants, H_2_O_2_ is mainly produced during the process of photosynthesis, respiration, photorespiration and wound/pathogen triggered ROS burst. Apart from the ROS burst which is mediated by membrane associated NADPH oxidases (rboh A∼F family) [10,12,44,45] under wounding and/or pathogen invasion, the photorespiration and electro transfer during photosynthesis constitute more than 90% of the H_2_O_2_ production [8,46] in healthy mesophyll cells. The former process is catalyzed by glycolate oxidase in peroxisome [47], whereas the latter is a non-enzymatic process (Mehler reaction) in chloroplast [48,49]. Both the photorespiration and electro transfer during photosynthesis require light, indicating the major H_2_O_2_ production is light dependent. Therefore, a high background level of H_2_O_2_ may exist in light grown plant, which interfere the observation of the H_2_O_2_ effects *in planta*.

To identify new motifs involved in H_2_O_2_ signalling, a preliminary survey on published microarray data which were collected from 5-d-old light grown seedlings (GSE5530 in NCBI GEO database) was conducted. However, only W-box and some well characterized light responsive motifs were identified (Supplementary Figure 1 online), suggesting the difficulty of using these data to identify new motifs. This is likely due to the high level of endogenous H_2_O_2_ in plant cells and exogenous H_2_O_2_ application most probably imposes oxidative stress. To reduce the endogenous H_2_O_2_ background, we grew *Arabidopsis* seedlings in the dark to eliminate light induced H_2_O_2_ production [36]. Exogenous H_2_O_2_ was then applied to the dark-grown seedling and transcriptome change was compared with those exposed to light. Interestingly, a substantial portion of genes in the genome were found to be co-regulated by H_2_O_2_ treatment and light, suggesting the H_2_O_2_ signal transduction occurs under light condition [36]. In addition, a GGGCC sequence motif (site II like) was identified and validated using gel shift assay. Genome wide pattern search indicated this motif tends to occur in the upstream regions adjacent to the transcription start site in higher eukaryotic species, such as *Mus musculus, Homo sapiens* and *Arabidopsis*, but not *Saccharomyces cerevisiae*. EMSA results confirmed this observation and the binding activity were found significantly increased under H_2_O_2_ treatment. C/D family small nucleolar ribonucleoprotein particles (snoRNPs) were identified as the binding nuclear factors (NFs) of site II motif, and the result was verified by shift-Western blotting and immunodepletion experiments.

**Figure 1.**
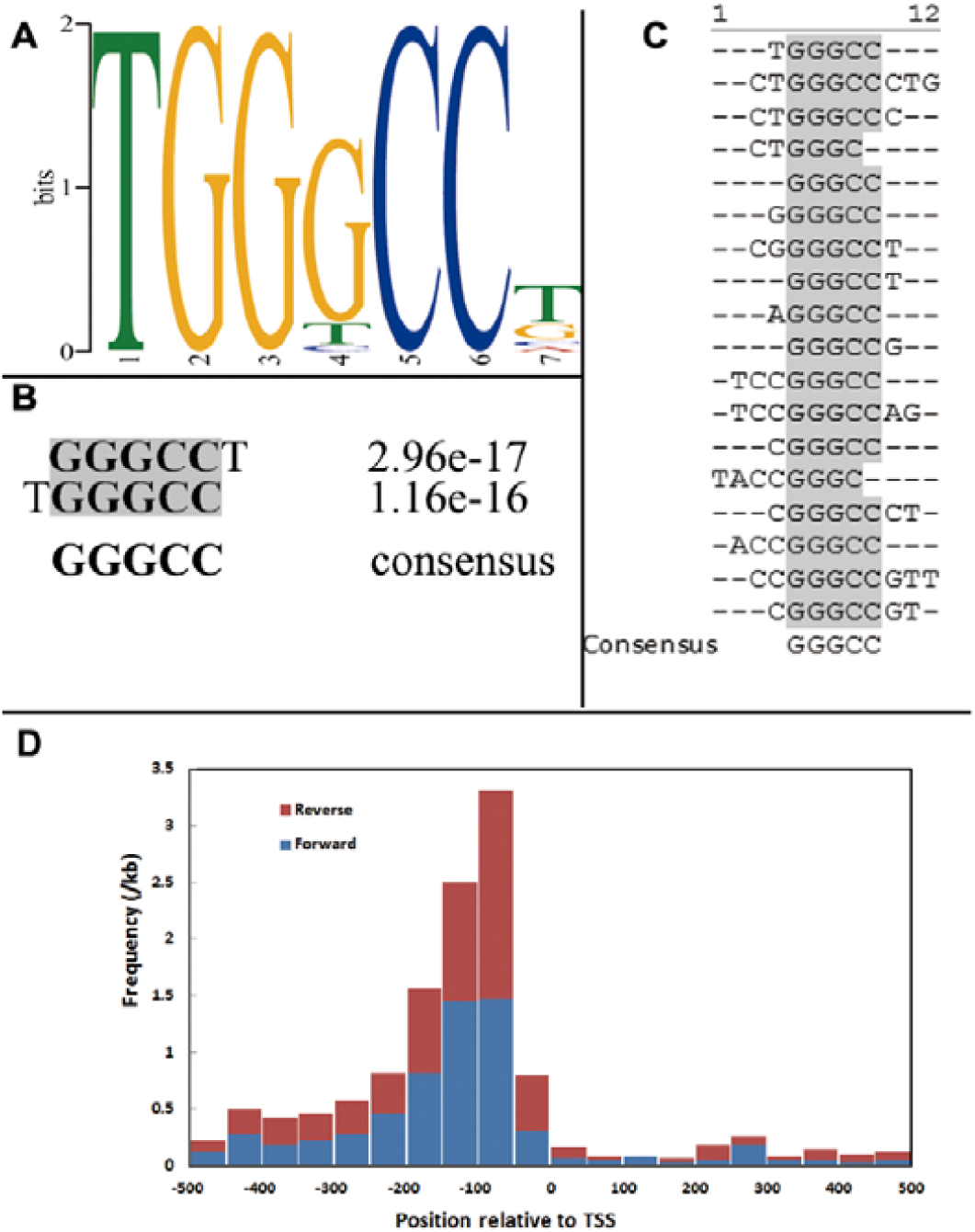
Computational identification of GGGCC motif from the upstream sequences of light and H2O2 co-upregulated genes. **A**. Sequence logo of the motif discovered by MEME software. **B & C**. Sequence alignments of the outputs with the highest score from TAIR Motif Analysis (B) and Weeder (C) software. D. GGGCC motif distribution frequencies in the −500bp region of the transcriptional start site (TSS) of light and H2O2 co-upregulated genes. Both the forward (in blue color) and reverse strands (in red color) were calculated.

## 2 Results

### 2.1 Identification of a site II like motif motif from the promoters of co-upregulated genes

To investigate if any common motif exists in the promoters of the 1021 light and H_2_O_2_ co-upregulated genes [36], we examined their upstream sequences up to −500 bp using MEME program [50] (E-value threshold of 0.001). A 7-nucleotide (nt) motif TGGGCC(T/G) was confidently identified (e-value of 1.2E-71) (Figure 1A). Subsequent search using TAIR Motif Analysis tool (http://www.arabidopsis.org/tools/bulk/motiffinder/index.jsp) and Weeder 1.42 [51] also identified the same motif (Figure 1B & C). The alignment result from the Weeder outputs suggested that the first and the last nucleotide are not always necessary (Figure 1C). The motif was therefore shortened to GGGCC. Because this motif is very similar to the previously reported site II motif (TGGGCC) in the promoter of rice and Arabidopsis PCNA gene [52,53], we therefore consider it as a more compact form of site II motif. By scanning the region from −500 bp to +500 bp around the transcription start site (TSS) in the 1021 co-upregulated genes, we found this site II like motif more frequently locates in the upstream region, especially from −50 to −200 bp (Figure 1D). The abundance of this motif in this region suggests its possible regulatory roles in H_2_O_2_ signalling.

### 2.2 Verification of the GGGCC Motif by Gel Mobility Shift Assay

To verify the nuclear factor (NF) binding activity of this site II like motif, a gel mobility shift assay (EMSA) was performed using the R-box (containing site II motif) as the probe (Figure 2A). Nuclear proteins were prepared from *Arabidopsis* PSB-D protoplast cells, and 5’-digoxigenin labelled probe (hot probe) was added to form NF-DNA complexes. The fish sperm DNA was included to suppress non-specific bindings. Non-labelled probes (cold probe) with or without mutations were used as competitors to ensure the binding is sequence specific (Figure 2A). As illustrated in Figure 2B (lane 2) and Figure 3A, at least 5 retarded bands were detected. These bands can be classified into 2 groups: fast group (NF F1, F2, F3) and slow group (NF S1 and S2), suggesting multiple NFs may participate in forming the NF-probe complexes. When unlabelled cold probe was added, the retarded bands signals were reduced (Figure 2B, lane 3, 4 and 5). The fact that 125 folds excessive cold probe almost completely blocked the NF S and the F group retarded bands (Figure 2B, lane 5) demonstrated that the NF-probe binding is R-box specific.

**Figure 2.**
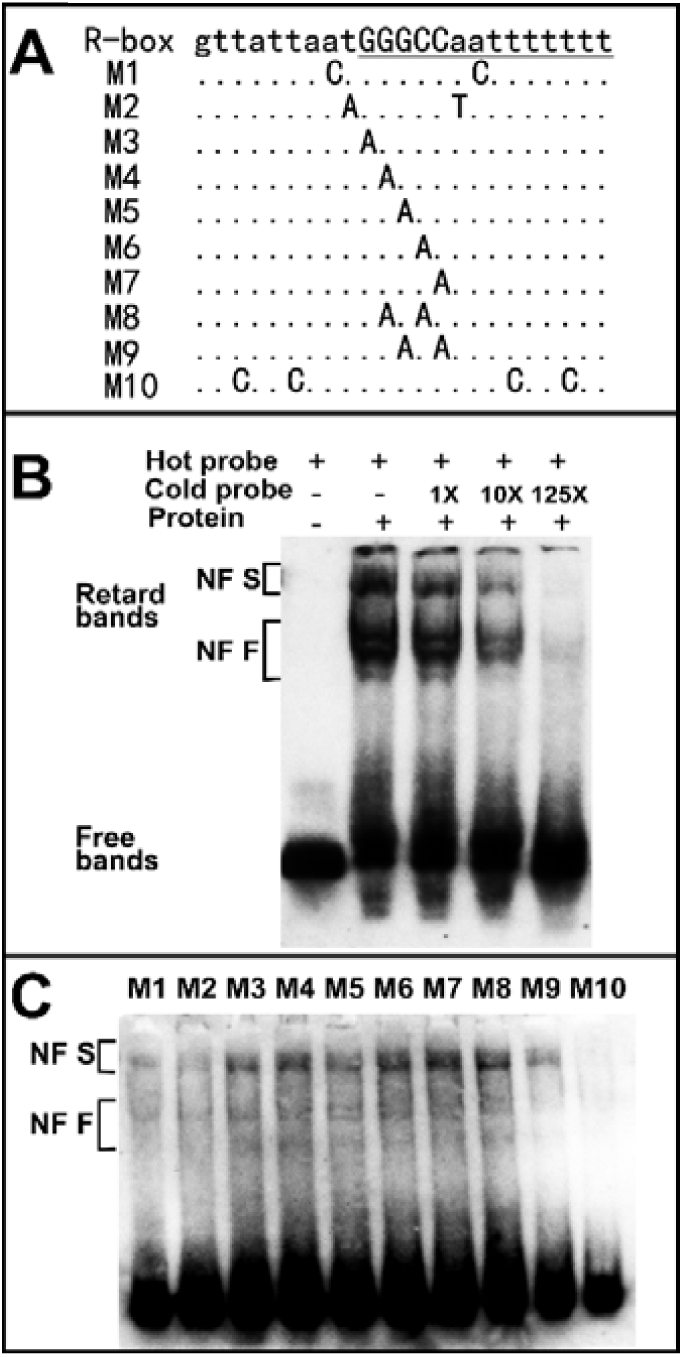
The validation of GGGCC motif with gel mobility shift assay. **A**. Sequences of the probes and the mutation sites. **B**. Gel mobility shift assay with R-box probe. Thirty microgram nuclear proteins (lane 2-4, left to right) from PSB-D protoplast cells were incubated with 1 ng 5’-Digoxigenin labeled R-box probe and then separated by PAGE. Probes were transferred to positive charged nyon membrane and signals were detected with Digoxigenin detection kit (Roche). A negative control was performed by replacing nuclear protein with H2O (lane 1). The non-labelled cold probe was used as competitor in the binding reaction. The signal bands of specific complexes and free non-binding probe were indicated as “retard bands” and “free bands” respectively. The NF F represents nuclear factor with fast mobility, and NF S represents nuclear factors with slow mobility. **C**. Gel mobility shift assay using mutated probes as competitors. 125 ng cold probes that mutated at indicated sites were bound with 30 μg *Arabidopsis* protoplast nuclear proteins for 10 min. Then 1ng labeled R-box probe was added and incubated for further 20 min. The reactions were separated by PAGE and bands were detected.

**Figure 3.**
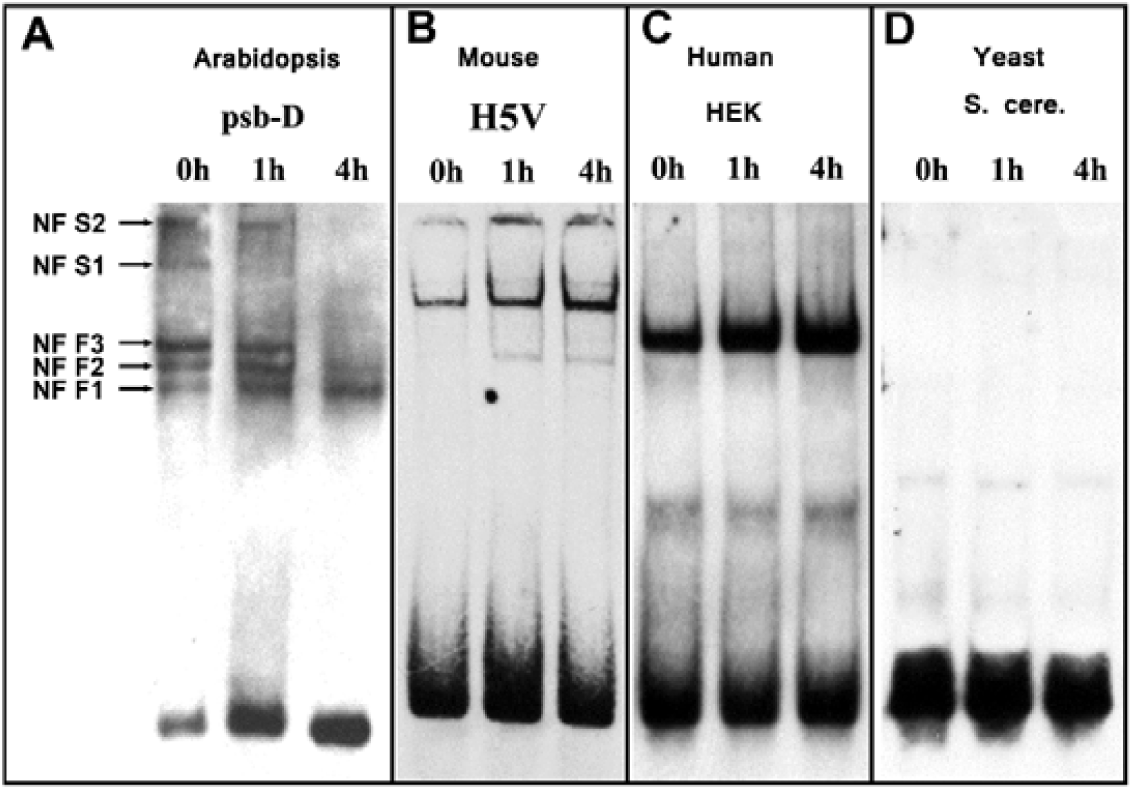
The site II like motif responds to H2O2 in Arabidopsis, mouse and human, but not yeast. The *Arabidopsis* suspension cells, mouse H5V cells, human HEK cells and *Saccharomyces cerevisiae* cells were treated with H2O2 for the indicated time and then the nuclear proteins were extracted. Twenty microgram nuclear proteins of each *Arabidopsis*, mouse and yeast extraction and five microgram human nuclear proteins were bound to 1 ng 5’-Digoxigenin labeled R-box probe.

To further examine if GGGCC is the core binding sequence, 125 folds mutated cold probes (Figure 2A, M1 to M10) were added as competitors. The mutations in the core sequence would impair the competitor ability of cold probes, so that the hot probe could reproduce retarded bands. As shown in Figure 2C, probes containing single nucleotide mutation or two mutations in the GGGCC sequence significantly restored the NF S bands, and the NF F bands also showed increased signals (Figure 2C, M3-M9). Two nucleotides mutations at the adjacent sites (M1 and M2) or four mutations at the distant sites (M10) did not give much signal restoration (Figure 2C). These data indicated that GGGCC is the core sequence for NF binding.

### 2.3 GGGCC motif is widely distributed in eukaryotic gene promoters

By searching the genome data, we found that the GGGCC motif widely exists in many eukaryotic species, such as *Saccharomyces cerevisiae, Mus musculus, Homo sapiens* and *Arabidopsis thaliana et. al*. These genomes were surveyed to calculate the GGGCC distribution frequencies in different sub-regions, including the upstream and downstream regions of genes, exons, introns, and the whole genome. Both forward and reverse strands were searched as GGGCC is not palindromic. We found that this motif is more frequently distributed in the upstream regions of genes in mouse, human and *Arabidopsis*, but not yeast genome (Table 1). The most frequent distribution was recorded within the −250bp regions for *Arabidopsis* (0.977 per kb), mouse (4.028 per kb) and human (5.974 per kb) genome. The high frequency of occurrence in this region suggests a possible regulatory role of this motif.

**Table 1.**
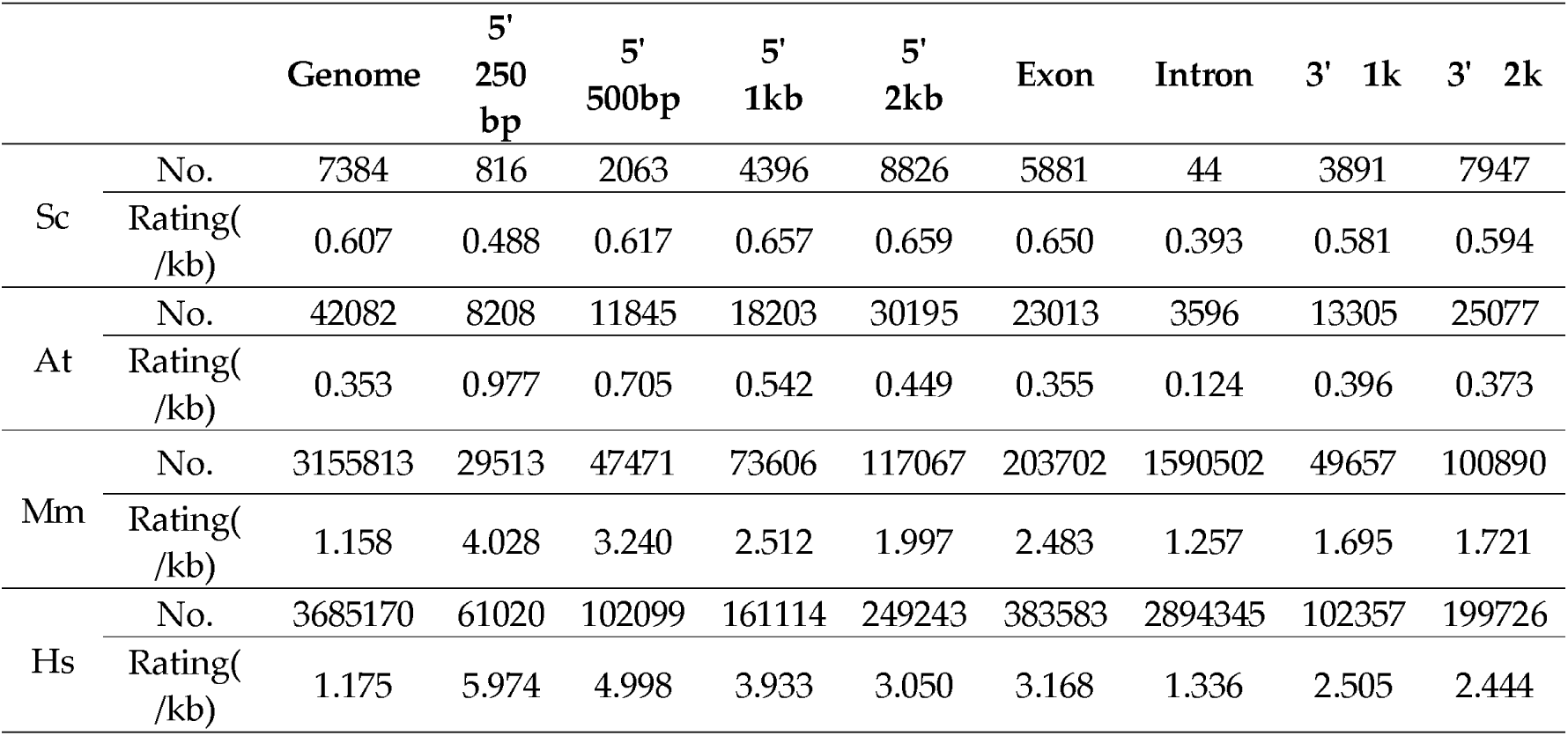
GGGCC distribution pattern in different genomes. GGGCC motif distribution frequencies in different subsets in the genomes of *Saccharomyces cerevisiae* (Sc), *Arabidopsis* (At), *Mus musculus* (Mm) and *Homo sapiens* (Hs). Motif frequencies were calculated on both forward and reverse strands in the upstream sequences up to 250bp, 500bp, 1kb and 2kb, exons, introns, downstream 1kb, 2kb, and whole genome using the annotated genomic data. The frequencies were given as the occurrence of the motif per kilobase pair in the specific regions.

It is also noteworthy that site II like motif is much highly rated in human and mouse genome compared to Arabidopsis. For mammalian species, the GGGCC frequencies within the −1kb region are 2.512 and 3.933 per kb for human and mouse respectively, i.e. roughly each gene contains 2.5 and 4 elements in this particular region. The frequency in the same region is much lower for Arabidopsis, which is only 0.542 per kb. In addition, the lack of enriched site II like motif in yeast genome implied that it may not function in this single cell eukaryotic species.

### 2.4 GGGCC motif is ROS responsive in higher eukaryotic species

To experimentally verify the site II like motif in yeast, mouse, human and Arabidopsis, nuclear proteins were extracted from the cells of each species and gel mobility shift assay was performed. The cell lines used for this test were: Arabidopsis PSB-D protoplast cells, human embryonic kidney 293 cells (HEK), mouse embryonic heart endothelial cells (H5V) and yeast Saccharomyces cerevisiae spheroplast cells. The cells were treated with either 1 mM (for yeast and animal cells) or 5 mM (for Arabidopsis cells) H_2_O_2_ to test if this site II like motif respond to ROS stress. As shown in Figure 3, nuclear proteins extracted from H5V and HEK cells also produced shifted binding bands (Figure 3B & C), which were proved to be site II like motif specific by competition experiments (Supplementary Figure 2 online). Interestingly, no band was observed for yeast (Figure 3D). These results comply with the motif distribution pattern, that is, the site II like motif is not enriched in the upstream region of genes in yeast genome (Table 1).

Nuclear proteins extracted from the H_2_O_2_ treated PSB-D protoplast, H5V and HEK cells showed altered motif binding activities (Figure 3A, B, & C), indicating this site II like motif was H_2_O_2_ responsive in these cells. For H5V and HEK cells, 1mM H_2_O_2_ treatment resulted in increased retarded bands signals within 1 hour, and the signal intensity continued to increase after 4 hours. For Arabidopsis protoplast cells, the retarded bands showed more complicated pattern upon H_2_O_2_ treatment (Figure 3A). NF F1 band signal was gradually increased after 1h and 4h treatment. The NF F2 band signal was increased within 1h, and was then decreased after 4h. The NF F3 and S1, S2 bands signal was decreased during H_2_O_2_ treatment, and could not be detected after 4 h. These patterns suggest the likely distinct regulatory roles of these NFs.

### 2.5 NOP56 and fibrillarin 2 proteins were Identified from NF+ samples

When our EMSA results were compared with those from the previously reported site II motif (TGGGCC) (Trémousaygue et al. 2003), we found that the binding profiles are different. To determine the identities of the binding proteins in our system, a purification procedure (Figure 4A) was adopted to scale up the nuclear proteins used for binding reaction. Totally 5 mg nuclear extractions were bound to 40 pmol biotin labelled R-box probes to form NF-probe complexes. The complexes were then recovered by the Streptavidin coupled magnetic beads and the NFs was eluted (NF+). Other 5 mg nuclear proteins were used as control experiment in parallel to recognise the unspecific bindings (NF-) by omitting probe in the reaction. The eluted NF+ and NF-was denatured and separated on SDS-PAGE gel. Totally 2 band groups with molecular weight of ∼67 kD and ∼31 kD respectively were successfully identified in the NF+ samples (Figure 4B). The bands were excised and in-gel digested with trypsin. The peptides were recovered and identified by ABI 4700 MALDI-TOF/TOF analyzer. The produced ion peaks were used to search against NCBInr database to find out the protein identities. The 31 kDa proteins was found to match the Arabidopsis fibrillarin 2 (FIB2) with a high score of 252. Totally 10 peptides were detected, which covered 36.9% of the AtFIB2 sequence (Table 2). The Arabidopsis genome contains 3 fibrillarins: AtFIB1 (AT5G52470), AtFIB2 (AT4G25630) and AtFIB3 (AT5G52490), which share high similarity in the conserved regions. Fib3 is probably a pseudogene (Barneche et al, J. Biol. Chem 2001).One of the identified peptides, GPAGRGGMK, were only found in the AtFIB2 sequence, and therefore fibrillarin 2 was deduced to be included in the binding complex. Even so, the FIB1 cannot be excluded, considering only 36 % of Fibrillarin protein was covered and the high similarity between FIB1 and FIB2 sequences.

**Table 2.**
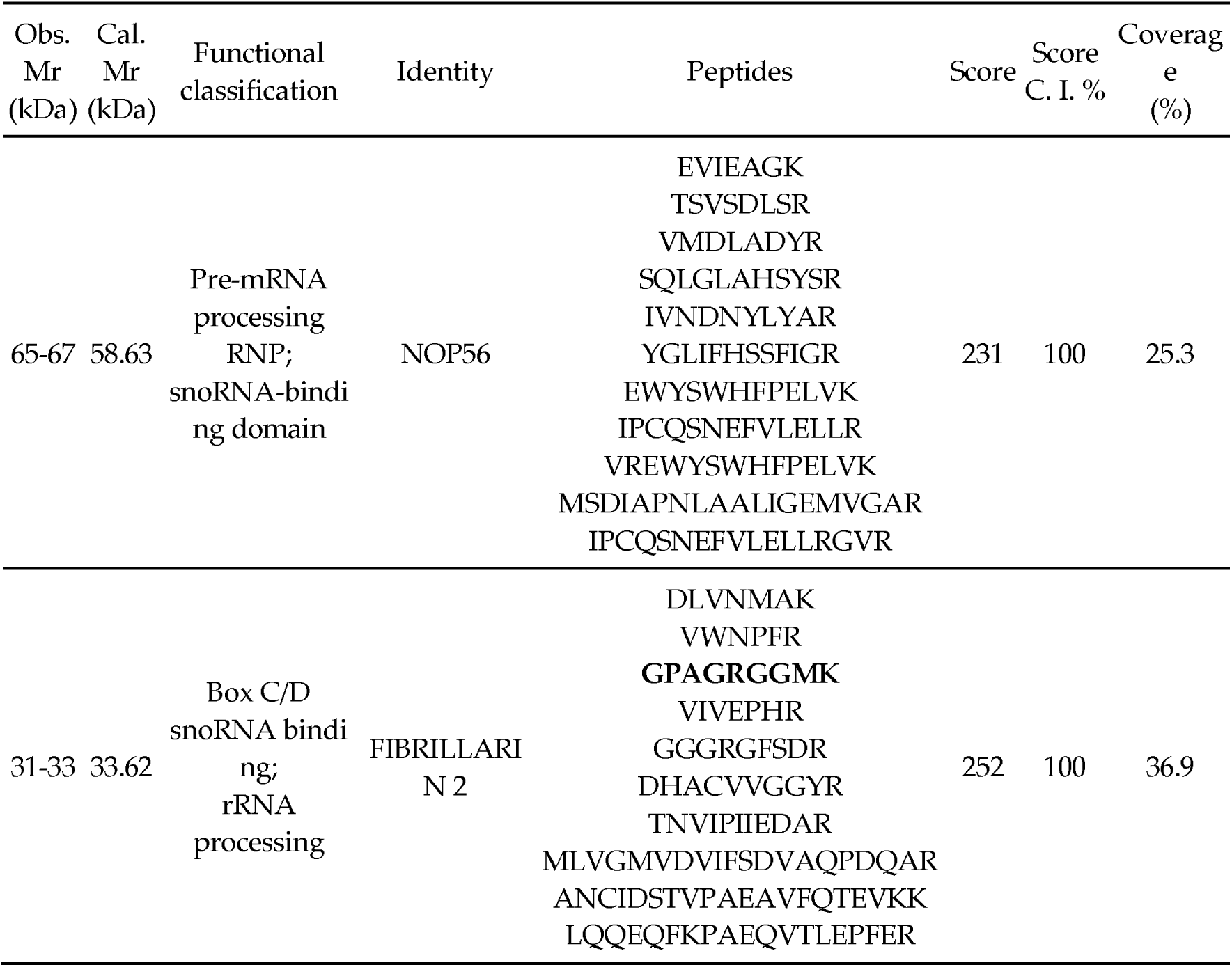
Peptides identified from the mixtures of digested target proteins. The protein bands were excised and in-gel digested as described in the Material and Methods. Mass spectrometric analyses were performed on a MALDI-TOF/TOF tandem mass spectrometer ABI 4700 proteomics analyzer (Applied Biosystems). All mass spectra acquired were internally calibrated with the ions of trypsin autolysis peptides and then searched against the NCBInr database. The matched peptides of each protein were listed. The candidates with the maximum number of matched peptides and a calculated Mr. value nearest to the observed value was chosen to report.

**Figure 4.**
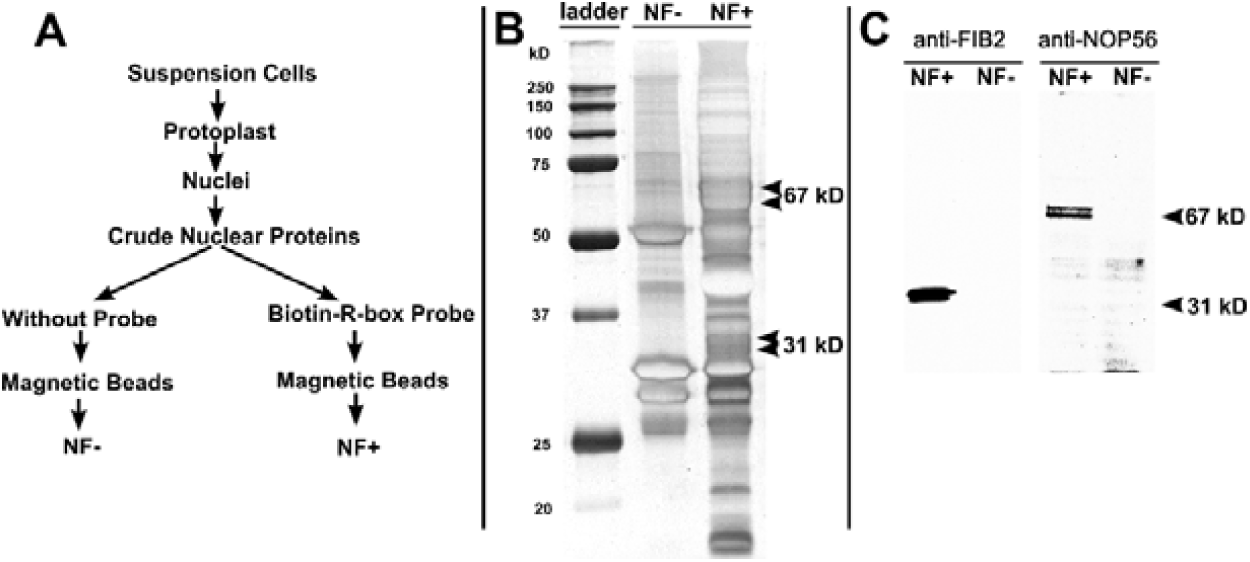
The identification of GGGCC motif binding proteins. **A**. Diagram showing the purification procedures of NF+ and NF-. **B**. SDS-PAGE of the NF+ and NF-proteins. NF+ and NF-were purified from each 5 mg nuclear proteins, denatured with SDS loading buffer, run on 12% SDS-PAGE, and silver stained. Arrow heads indicate the protein bands that only found in NF+. **C**. Western blotting to validate the MALDI-TOF-TOF identified protein candidates. 12 μL of NF+ and NF-were mixed with 3 μL 5X SDS loading buffer, boiled and separated on 12% SDS-PAGE gel. Then the proteins were transferred onto nitrocellulose membrane and blotted with 1:500 dilution of antibody anti-fibrillarin or anti-NOP56.

As for the 67 kDa bands, in total 11 peptides were detected and matched to NOP56 protein with a score of 231. Two NOP56 homologs (AT3G12860 and At1g56110) which share more than 97% peptide similarity were found in *Arabidopsis* genome. All the detected 11 peptides can be found in the peptide sequences of these two genes, suggesting the peptides are probably derived from both NOP56 isoforms.

The NF+ and NF-were denatured by SDS-loading buffer and separated on SDS-PAGE gel. Antibody anti-FIB2 and anti-NOP56 were used to detect the corresponding target proteins. The Western blotting results confirmed the identity of 31 kDa FIB2 protein and the 67 kDa NOP56 protein.

Interestingly, both the identified proteins are snoRNP proteins with snoRNA binding and ribosomal RNA processing function (Table 2). Searching the *Arabidopsis thaliana* Protein Interaction Network (AtPIN) database [54] revealed that NOP56 and FIB2 are interactive proteins, supporting our hypothesis that AtFIB2 and AtNOP56 proteins are components of the NFs which bind with the site II like motif.

Immunodepletion was performed to further validate AtFIB2 and AtNOP56 are indeed involved in site II like motif binding. The nuclear proteins were subjected to immunodepletion with antibody anti-fibrillarin 2 and anti-NOP56. The depletion of fibrillarin 2 significantly reduced R-box probe binding activity when compared with those not depleted or depleted with pre-immune mouse serum. However, depletion of NOP56 did not result in much reduction in the R-box probe binding activity (Figure 5A), likely due to the fact that the antibodies were produced with short conserved peptides and therefore may have low reactivity with the native proteins. We then chose shift-Western [55] for *in situ* detection of target proteins in the retarded bands. As shown in Figure 5B, FIB2 was detected in the bands corresponding to both NF F and S group EMSA shifted bands. NOP56 protein was only found in the location corresponds to NF S1, S2 and F3 EMSA shifted bands. The successful detection of FIB2 and NOP56 in the NF-probe complexes demonstrates that these candidate proteins are involved in the formation of nuclear factors that bind to the R-box probe.

**Figure 5.**
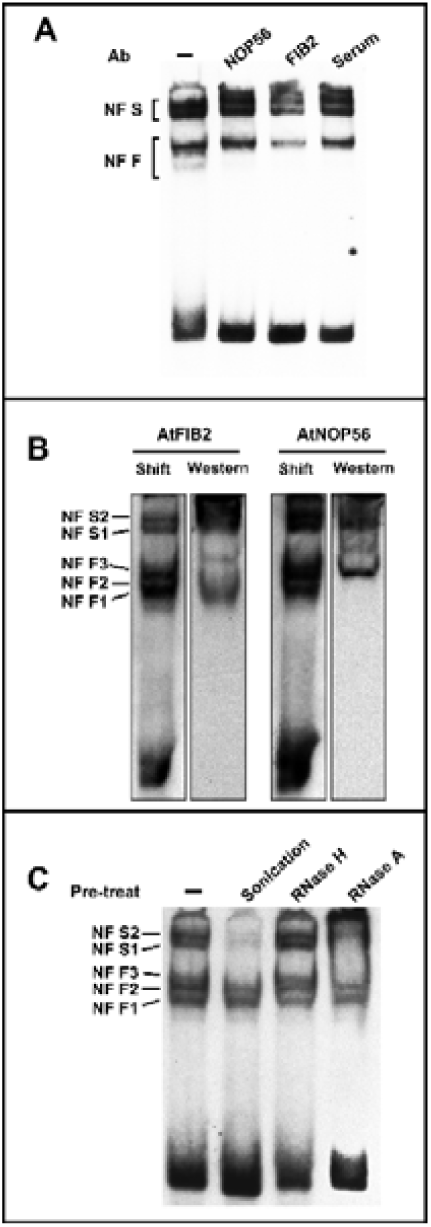
FIB2 and NOP56 proteins, and RNA were detected in R-box probe binding NFs. **A**. Binding activity detection of the nuclear proteins depleted with antibody anti-fibrillarin and anti-NOP56, or pre-immune serum. **B**. Shift-Western blotting for detecting FIB2 and NOP56 in the NF-probe complexes. Nuclear proteins were bound with R-box probe and separated on 6% PAGE gel. Then the blot was transferred onto stacked nitrocellulose membrane and positive charged nylon membrane. The probe was detected from nylon membrane and Western blotting was performed to detect the target protein from nitrocellulose membrane with indicated antibodies. Shift, probe signals detected from nylon membrane; Western, indicated protein signals detected from nitrocellulose membrane. **C**. Binding activity of the nuclear proteins pre-treated with sonication, RNase H and RNase A. 30 μg nuclear proteins were pre-treated with sonication, 5U RNase H or 5U RNase A for 10 min at 25 °C, then bound with 1 ng R-box probe for binding activity detection.

### 2.6 Molecular cloning and detection of box C/D family snoRNA from NF+ samples

Because fibrillarin and NOP56 are core proteins associated with box C/D snoRNA [56], the identification of FIB2 and NOP56 from NF+ suggests that box C/D family snoRNPs complexes may be involved in site II like motif binding. To test this hypothesis, nuclear extract was pre-treated with RNase H and RNase A to deplete small RNA. RNase A can degrade both single- and double-stranded RNA while RNase H only degrades RNA in DNA-RNA hybrids. As shown in Figure 5C, 10 min incubation with RNase A removed NF S1 and F3 bands, and NF S2, F1 and F2 bands signals were significantly reduced. The incubation with RNase H did not show much signal reduction compared with the control. The RNase depletion results indicate that RNAs participate in the binding with R-box probe. Interestingly, we found that 10 min sonication also reduced the NF S1, S2 and F3 band signals, suggesting these NFs are not tightly associated.

To identify the RNA species involved in the site II like motif binding, small RNA was extracted from the NF+ elution and used to construct a cDNA library. In total about 100 colonies were picked up and sequencing results showed that 30 sequences belong to 18 snoRNA species (14 C/D family and 4 H/ACA family)(Supplementary Table 2 online). To verify these snoRNAs are specific binding, RT-PCR was used to examine the abundance of each snoRNA in the NF+ and NF-samples using gene specific primers. In total, 3 C/D family snoRNAs (R63, U24 and Z15) were detected only in the NF+ sample, and the products are of exactly the same size with the corresponding bands from whole cell RNA (Figure 6A, B & C). No corresponding bands were detected from either NF-fraction or NF+ fraction that was not reversely transcribed, demonstrating R63, U24 and Z15 snoRNAs are specifically enriched in the NF+ fraction, and are the components of NFs that bind with R-box probe.

**Figure 6.**
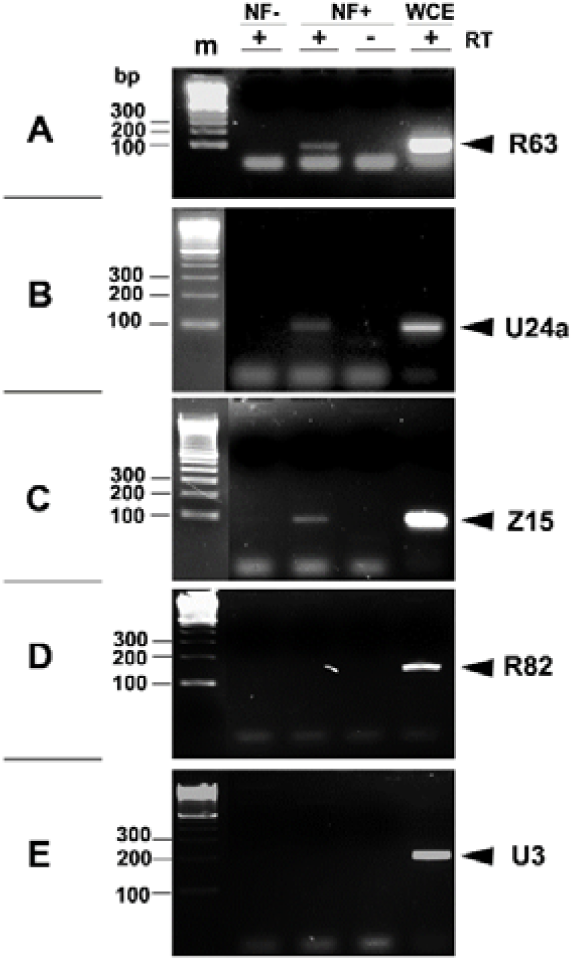
Identification of snoRNAs in NF+ by RT-PCR. Specific primers were designed to amplify R63, U24a, Z15, R82 and U3 based on the sequencing results and *Arabidopsis* gene sequences in the Genbank. RNAs extracted from NF-, NF+ and whole cell extracts (WCE) were reverse transcribed (RT) with random primers. RT-PCR was carried out using corresponding primer pairs and individual cDNA samples. NF+ RNA was used as RT negative control to monitor the possible amplification from contaminated genome DNA. The amplification products of WCE cDNA was used to indicate the size of each snoRNAs.

## 3 Discussion

ROS has great advantages to function as signals, e.g. the dynamic ROS levels across tissues and organs, quick conversion between different forms, fast long-distance propagation, easy metabolism and transport in cells et al [57]. The production of ROS can be triggered by many biotic and/or abiotic factors such as pathogens, wounding, heat and cold shocks et al [58–61]. In addition, ROS are produced by diverse mechanisms in specific compartments in the cells. The question is how the generated ROS can be recognized by the cell as specific signals to activate the gene expression accordingly in order to adapt to the external cues. Currently three possible mechanisms regarding the specificity of ROS signal pathway [23] have been proposed, and in each case a universal ROS signal transduction is required because ROS generation is universal in higher species.

ROS signal transduction requires both cis-elements and transcription factors that bind to the regulatory elements. However, up to now only a few motifs were experimentally verified to be ROS responsive in plants [62]. A genome wide *in silico* prediction have identified a NRXe2 motif with a cognate sequence of TGACGTCA, which specifically responds to H_2_O_2_ but not ABA or sucrose [41]. Another 28 bp conserved CORE motif was found to respond to oxidative stress in rice [40]. Other motifs such as tobacco activation sequence-1 (as-1)-like element [63], antioxidant-responsive element in maize [39], O3-responsive region of grapevine stilbene synthase [38] were also oxidative stress related. These studies extended our understandings about ROS signalling network in plants. In this study, we identified a ROS responsive motif from *Arabidopsis* by searching the upstream sequences of the genes co-upregulated by both H_2_O_2_ and light [36]. This motif contains a core sequence of GGGCC, which was previously identified from the promoters of phytochrome A induced genes in *Arabidopsis* [64]. However, because H_2_O_2_ responsive genes highly overlap with light responsive ones and ROS signalling is inevitable under light condition, it is difficult to distinguish whether the GGGCC motif is light induced or ROS induced [36]. On the other hand, considering the facts that R-box probe binding activity was increased under H_2_O_2_ treatment in both plant and animal cells (Figure. 5), it is more reasonable to deduce that GGGCC is “ROS induced” rather than “light induced” because no light was imposed on these cells in our system. We also noted that GGGCC motif is very similar to the previously identified site II motif (TGGGCC) in the promoters of plant *proliferating cellular nuclear antigen* (*PCNA*) gene [52,65] and nuclear genes encoding components of the oxidative phosphorylation (OxPhos) machinery [66], which has been demonstrated to regulate gene expression in cycling cells in root primordia by binding with bHLH transcription factors (PCF1 and PCF2) [52,53]. The site II motif (TGGGCC) is also over-represented in the promoters of cold- and dehydration-induced genes [67]. Although GGGCC is considered as a compact form of site II motif (site II like), some differences still exist when compared with previously reported site II motif. First, as demonstrated in Figure 2C, T(GGGCC) is not essential for R-box probe to bind with NFs. Second, R-box probe and previously identified site II motif bind to different NF complexes as indicated by EMSA experiments (Figure 2, 3 & 5) [53]. Site II motif specifically bind to bHLH family transcriptional factors [52,53], while R-box probe is associated with C/D family snoRNP complexes.

The identification of site II like motif binding NFs revealed that the box C/D family snoRNAs are the components of the complexes bind with this motif. SnoRNA is small nucleolar non-coding RNA that participates in post-transcriptional RNA modifications, in which pseudouridylation and 2’-O-ribose methylation are the major types. Two classes of snoRNAs universally exist in eukaryotes in the forms of small nucleolar ribonucleoprotein (snoRNPs): C/D family and H/ACA family. H/ACA class snoRNP directs isomerization of uridine to pseudouridine (ψ) while C/D class snoRNP guides site-specific 2’-O-ribose methylation [68]. Both the C/D and H/ACA snoRNAs associate with core proteins which carry out enzymatic catalyzation for the RNA modifications. Fibrillarin, NOP56, NOP58 and 15.5-kDa are the four core proteins of C/D snoRNPs [69]. Fibrillarin is a ribose-2’-O-methylase that catalyzes 2’-O-ribose methylation, while NOP56 is required for maintaining the stability of snoRNAs. The identification of fibrillarin 2 and NOP56 proteins from the NF+ fraction strongly suggests that C/D class snoRNP is associated with the site II like motif. A cDNA library was constructed using the RNA extracted from NF+ fraction, and 18 snoRNA species were identified. Three C/D family snoRNAs, including R63, U24 and Z15, were further validated in the NF+ fraction by RT-PCR.

It has been reported that motifs in the 5’ external transcribed spacer (ETS) of rRNA genes can bind with C/D family snoRNPs, which is a pre-rRNA processing complex that assembles on the rDNA prior to binding to the nascent transcript [70,71]. In this study, we found C/D snoRNPs bind with the ROS responsive site II like motif, which is highly abundant in the upstream regulatory sequences. The targets spectrum of box C/D snoRNAs are expanding, e.g. they are known to be involved in spliceosomal snRNA [72,73] and tRNA [74] modification, pre-mRNA alternative splicing [75–77], and rRNA 2’-O-ribose methylation. However, it is still quite unusual for the C/D snoRNA to bind to the cis-elements. Because the −50 bp to −250 bp upstream region is most important for gene expression regulation, the molecular function of the GGGCC motif binding snoRNAs is of great interest. In yeast, fibrillarin homolog Nop1p protein associates with C/D snoRNA coding genes and functions as *trans*-acting factor responsible for box C/D snoRNA 3’-end formation by preventing read-through to snoRNA downstream gene by RNA polymerase II [78]. The identification of box C/D snoRNA associated proteins fibrillarin and NOP56 bound with site II like motif led us to hypothesize that snoRNP may regulate the function of polymerase II, which catalyzes the transcription of pre-mRNA. We also found that the GGGCC in the −500bp upstream region is significantly associated with alternative splicing event for *Arabidopsis* genes (unpublished data). Therefore, we deduce it is also possible that this site II like motif may provide an anchor site for box C/D snoRNPs, which are involved in the regulation of alternative splicing [75–77]. Further experiments are being carried out to test this hypothesis.

ROS signalling widely exists in prokaryotic and eukaryotic organisms such as bacteria, yeast, animals and plants. Recently, an LRR receptor kinase HPCA1 was identified as H_2_O_2_ in Arabidopsis [37]. However, no common regulatory pathway has been identified so far. In addition, no universal regulon has been found in plants and animals. Our study identified a common ROS responsive motif exists in both plants and animals by computational prediction and experimental validation, suggesting the common signalling pathway in response to ROS cross kingdoms. Considering the wide distribution of GGGCC among genomes and its quick response to H_2_O_2_, we also speculate this motif may function at the early steps in the ROS signalling pathway.

## 4 Materials and Methods

### 4.1 Plant materials and culture conditions

The suspension cell line PSB-D (TAIR Stock CCL84840) was a gift from Prof. Jiang Liwen at CUHK. The cell line is routinely subcultured every 5 days in the dark as reported [79]. The H5V murine heart capillary endothelial cells and human embryonic kidney HEK-293 cells were cultured in DMEM medium enriched with 2 mM l-glutamine, 10% heat-inactivated fetal bovine serum (FBS), 1% penicillin /streptomycin (Gibco, Life technology) at 37 °C under controlled conditions (95% O_2_ and 5% CO_2_). The *Saccharomyces cerevisiae* was inoculated in YPD medium and cultured at 28 °C with shaking (225 rpm).

For H_2_O_2_ treatment, a final 5mM H_2_O_2_ was added to PSB-D protoplast cells. For animal and yeast cells, the H_2_O_2_ concentration was 1mM. After 1h and 4h treatment, the cells were harvested and used for nuclear proteins extraction.

### 4.2 Nuclear protein extraction

*Arabidopsis* protoplast [79] and yeast spheroplast [80] was isolated as described, and used for nuclear protein extraction according to the reported method with modifications[81]. In short, the animal cells, plant protoplast cells or yeast spheroplast cells were suspended in nuclear isolation buffer (10 mM HEPES, pH 7.6, 5 mM MgCl2, 800 mM sucrose, 5 mM EDTA, 1 mM dithiothreitol and 1 mM phenylmethylsulfonyl fluoride, 1% protease inhibitor cocktail (Sigma, St. Louis, USA)) and vortexed vigorously for 1 min, then centrifuged at 12 000 g for 5 min and the supernatant was discarded. The precipitated nuclei were re-suspended in nuclear isolation buffer and washed twice. After carefully pipetting off all the liquid, the crude nuclei were frozen in liquid nitrogen and thawed on ice, then suspended with low salt buffer (20 mM HEPES, pH 7.6, 25% (v/v) glycerol, 20 mM KCl, 1.5 mM MgCl2, 0.2 mM EDTA, 0.5 mM dithiothreitol, 0.1% (v/v) nonidet P-40, 1 mM phenylmethylsulfonyl fluoride and 1% protease inhibitor cocktail) and incubated on ice for 5 min. After that, 4 M KCl was added in small aliquots to a final concentration of 0.5 M. The nuclei were incubated in cold room with agitation for 30 min and then centrifuged at 16 000 g for 30 min. The supernatant was carefully transferred to a new tube and stored at −80°C.

### 4.3 Gel mobility shift assay

The sequences of the double-stranded oligonucleotides used for the gel mobility shift assay were R-box probe (5’-GTTATTAATGGGCCAATTTTTTT-3’), M1 (5’-GTTATTACTGGGCCACTTTTTTT-3’), M2 (5’-GTTATTAAAGGGCCTATTTTTTT-3’), M3 (5’-GTTATTAATAGGCCAATTTTTTT-3’), M4 (5’-GTTATTAATGAGCCAATTTTTTT-3’), M5 (5’-GTTATTAATGGACCAATTTTTTT-3’), M6 (5’-GTTATTAATGGGACAATTTTTTT-3’), M7 (5’-GTTATTAATGGGCAAATTTTTTT-3), M8 (5’-GTTATTAATGAGACAATTTTTTTT-3’), M9 (5’-GTTATTAATGGACAAATTTTTTTT-3’) and M10 (5’-GTCATCAATGGGCCAATTCTTCTT-3’).

The 5’ Digoxigenin labeled R-box and non-labeled oligonucleotides were synthesized (Takara, Dalian, China). The binding mixture contained nuclear extracts, and 0.5 μg double strand fish sperm DNA (Roche, USA). Cold competitor probe was used as 125-fold molar excess or as indicated elsewhere. The binding buffer consist of 10 mM Tris-HCl, pH 7.5, 100 mM KCl, 10% (v/v) glycerol, 1 mM dithiothreitol, 0.1% (v/v) nonidet P-40 and 1 mM spermidine. The mixture was incubated for 10 min at 25°C, then 1 ng labelled probe was added. After further 20 min incubation, the reaction was loaded on 6% polyacrylamide gels containing 44.5 mM Tris, 44.5 mM borate, 1 mM EDTA, and 5% glycerol.

The blot was semi-dry transferred to Amersham Hybond™-N+ nylon membrane (GE Healthcare) and cross-linked under 0.12 J/cm2 UV irradiation. Digoxigenin was detected using DIG Nucleic Acid Detection Kit (Roche, USA).

### 4.4 Purification and identification of R-box probe binding NF proteins

Forty pmol biotin labelled R-box probe were bound with 5 mg Arabidopsis nuclear extracts for 20 min to form probe-NF complexes, then coupled to 1 mg Pierce® Streptavidin Magnetic Beads (Thermo Scientific, USA) and recovered with a MagnaBind™ Magnet. After 6 washes with binding buffer, the beads were eluted using 30 μL elution buffer (0.5 N NH4OH, 0.5 mM EDTA). The elution was regarded as NF+. A negative control (NF-) experiment was performed in parallel with 5 mg nuclear extracts with no probe. The NF+ and NF-samples were freeze dried, re-suspended in SDS-PAGE loading buffer, boiled and run on 12% SDS-PAGE gel.

The gel was silver stained, and the bands specifically found in NF+ samples were excised and distained. In-gel digestion and mass spectrometry (MS) analysis were performed as described previously [82,83]. Both MS and MS/MS data were searched against NCBInr database by Mascot search engine version 2.1 (Matrix Science, London, U.K.) in the GPS Explorer software v3.5 (Applied Biosystems, Foster City, CA) with the following parameters: taxonomy, *Arabidopsis thaliana*; enzyme, trypsin; allowed missed cleavages, 1; fixed modification, carbamidomethyl (C); variable modifications, oxidation; peptide tolerance, 0.5 Da; MS/MS tolerance, 0.5 Da; peptide charge, (+1); instrument, MALDI-TOF/TOF. Protein matches in MS/MS identification were considered valid when the Protein Score C.I. % is over than 95.

### 4.5 Antibodies

Anti-fibrillarin antibody [38F3] was from Abcam. Nop56 (Q-24) antibody sc-133839 was from Santa Cruz biotechnology, Inc. These antibodies have wide cross reactivity in plants and animals due to high homology in the immunogen peptide regions.

### 4.6 Western blotting

Twelve μL NF+ and NF-elutions were mixed with 3 μL 5X SDS loading buffer and boiled for 5 min, then subjected to SDS-12% PAGE. After that, the blot was transferred onto nitrocellulose membrane (Bio-rad) according to manufacturer’s instructions, then blocked in Tris-buffered saline with 0.1% Tween 20 (TBS-T) and 5% skim milk powder for 60 min. Primary antibodies were applied at a dilution of 1:500, and chemiluminescence protein detection was done as described with anti-mouse or anti-rabbit antibodies conjugated with horseradish peroxidase (HRP) from Cell Signaling Corp. A Promega ECL Western Blotting Substrate kit (W1001) was used for chemiluminescence enhancement.

### 4.7 Depletion tests

For immunodepletion of nuclear proteins, 1 μg fibrillarin 2 or NOP56 antibody was coupled to 30 μL protein-A agarose beads (Sigma) according to manufacturer’s instructions. Then the agarose beads were blocked with EMSA binding buffer containing 0.5 mg/mL bovine serum albumin (BSA) for 30 min at room temperature. Forty μg nuclear extracts were incubated with 15 μL of each pretreated protein-A agarose beads for 2h at 4 °C with agitation. The beads were removed by centrifuge and the supernatant was used to test R-box probe binding activity.

For sonication treatment, 19 μL binding mixture containing 4 μL nuclear extracts was sonicated using Branson Ultrasonic Cleaner 2510E-DTH (Branson Ultrasonic Corporation, USA) for 10 min at room temperature. For RNase depletion, 19 μL binding mixture containing 4 μL nuclear extracts and 5U Rnase H (NEB) or bovine pancreatic RNase A (Qiagen) was incubated for 10 min at 25 °C. After pre-treatment, R-box probe binding activity was detected by adding 1 ng hot probe.

### 4.8 Shift-Western assay

Shift-Western assay was performed based on Demczuk [55] et al. The EMSA assay was performed as described above. After separated on PAGE gel, the blot was transferred to stacked nitrocellulose membrane (Bio-rad) and positively charged nylon membrane (GE Healthcare) sequentially in cold Towbin’s buffer at 150 mA for 3h. The EMSA signal was detected on nylon membrane as described previously. Protein was detected by Western blotting as previously described.

### 4.9 Cloning and detection of snoRNAs

Small RNA was extracted from equal volume of NF+ and NF-samples (30 μL each) using Trizol reagents (Invitrogen). After digestion with DNase I, the RNA was precipitated and re-dissolved in 20 μL RNase free water. 10 μL each small RNA was used for cloning with a GeneRace® kit (Life Technologies) following the producer’s instructions with modifications. The small RNA was 3’ poly(A) tailed with Ambion® Poly(A) Tailing Kit (Life Technologies) and 5’ de-capped by using tobacco acid pyrophosphatase (TAP). The RNA was then ligated to GeneRace® RNA Oligo and reverse transcribed using a modified GeneRacer® Oligo dT primer (5’-GCTGTCAACGATACGCTACGTAACGGCATGACAGTG(T)_23_V-3’) and SuperScript® III reverse transcriptase. The cDNA was amplified with GeneRacer® 5’ and 3’ Nested Primers. The NF-sample was used to monitor the amplification cycles and to control non-specific products. The PCR were performed for 25 cycles and the products were separated on 3% agarose gel. 100-200 bp cDNA fragments abundant in NF+ sample were excised, recovered and cloned with TOPO TA Cloning® kit (Invitrogen). The colonies were picked up and sequenced to identify small RNA species.

RT-PCR was used to validate snoRNA. 10 μL DNase I digested small RNA extracted from NF- and NF+ were reverse transcribed with random primers and the cDNA was used for PCR by using GoTaq flexi DNA polymerase (Promega). 100 ng total RNA extracted from *Arabidopsis* protoplast cells were reverse transcribed and used as positive control to indicate the proper size of amplified products. Another RT reaction of 10 μL NF+ RNA with reverse transcriptase omitted was used as negative control. RT-PCR was run for 27 cycles with gene specific primers of each snoRNA. The primer sequences were listed in Supplementary Table 1 online.

### 4.10 Bioinformatics analysis

Bulk data were downloaded using Regulatory Sequence Analysis Tools (http://rsat.ulb.ac.be/rsat/). Genome data were extracted from UCSC Genome Browser database [84]. For motif discovery, up to 500 bp sequences upstream of selected genes were analyzed with MEME Suite software [50] and compared against a background file consisting of heptamer frequencies in the region of 22810 *Arabidopsis genes*.

## Supplementary Materials

Supplementary materials can be found online: Supplemental Tables 1-2, Supplemental Figures 1-2.

## Author Contributions

Project design: H.C. and D.G.; Experiment: H.C.; Computation: H.C. and M.S.; Manuscript writing: H.C. and D.G.

## Acknowledgments

We thank Dr. Fan Long for his help in analyzing the motif distribution in the genomes. We also thank Prof. Jiang Liwen at Chinese University of Hong Kong for providing the *Arabidopsis* PSB-D suspension cells and Prof. Yao Xiaoqiang for providing the H5V and HEK-293 cells.

## Abbreviations

H2O2: hydrogen peroxide
ROS: reactive oxygen species
NFs: nuclear factors
EMSA: electrophoresis mobility shift assay
snoRNP: small nucleolar ribonucleoprotein particles

## References

1. Malmstroem, B.G. Cytochrome c oxidase as a redox-linked proton pump. Chem. Rev. 1990, 90, 1247–1260, doi: 10.1021/cr00105a008.

2. Hammond-Kosack, K.E.; Jones, JDG. Resistance Gene-Dependent Plant Defense Responses. The Plant Cell 1996, 8, 1773–1791, doi: 10.1105/tpc.8.10.1773.

3. Mittler, R. ROS Are Good. Trends Plant Sci. 2017, 22, 11–19, doi: 10.1016/j.tplants.2016.08.002.

4. McCord, J.M.; Fridovich, I. Superoxide dismutase: The first twenty years (1968–1988). Free Radical Biology and Medicine 1988, 5, 363–369, doi: 10.1016/0891-5849(88)90109-8.

5. Waszczak, C.; Carmody, M.; Kangasjärvi, J. Reactive Oxygen Species in Plant Signaling. Annu Rev Plant Biol 2018, 69, 209–236, doi: 10.1146/annurev-arplant-042817-040322.

6. Bielski, B.H.; Arudi, R.L.; Sutherland, M.W. A study of the reactivity of HO2/O2-with unsaturated fatty acids. Journal of Biological Chemistry 1983, 258, 4759–4761.

7. Valavanidis, A. Generation of hydroxyl radicals by urban suspended particulate air matter. The role of iron ions. Atmospheric Environment 2000, 34, 2379–2386, doi: 10.1016/S1352-2310(99)00435-5.

8. Foyer, C.H.; Noctor, G. Redox sensing and signalling associated with reactive oxygen in chloroplasts, peroxisomes and mitochondria. Physiol Plant 2003, 119, 355–364, doi: 10.1034/j.1399-3054.2003.00223.x.

9. Neill, S.; Desikan, R.; Hancock, J. Hydrogen peroxide signalling. Current Opinion in Plant Biology 2002, 5, 388–395, doi: 10.1016/S1369-5266(02)00282-0.

10. Foreman, J.; Demidchik, V.; Bothwell, J.H.F.; Mylona, P.; Miedema, H.; Torres, M.A.; Linstead, P.; Costa, S.; Brownlee, C.; Jones, J.D.G.; et al. Reactive oxygen species produced by NADPH oxidase regulate plant cell growth. Nature 2003, 422, 442–446, doi: 10.1038/nature01485.

11. Cheeseman, J.M. Hydrogen Peroxide and Plant StressL: A Challenging Relationship. In Plant Stress; 2007; Vol. 1, pp. 4–15.

12. Miller, G.; Schlauch, K.; Tam, R.; Cortes, D.; Torres, M.A.; Shulaev, V.; Dangl, J.L.; Mittler, R. The Plant NADPH Oxidase RBOHD Mediates Rapid Systemic Signaling in Response to Diverse Stimuli. Sci. Signal. 2009, 2, ra45, doi: 10.1126/scisignal.2000448.

13. Frugoli, J.A.; Zhong, H.H.; Nuccio, M.L.; McCourt, P.; McPeek, M.A.; Thomas, T.L.; McClung, C.R. Catalase Is Encoded by a Multigene Family in Arabidopsis thaliana (L.) Heynh. Plant Physiol. 1996, 112, 327–336, doi: 10.1104/pp.112.1.327.

14. Chelikani, P.; Fita, I.; Loewen, P.C. Diversity of structures and properties among catalases. Cellular and Molecular Life Sciences (CMLS) 2004, 61, 192–208, doi: 10.1007/s00018-003-3206-5.

15. Mhamdi, A.; Queval, G.; Chaouch, S.; Vanderauwera, S.; Van Breusegem, F.; Noctor, G. Catalase function in plants: a focus on Arabidopsis mutants as stress-mimic models. J. Exp. Bot. 2010, 61, 4197–4220, doi: 10.1093/jxb/erq282.

16. Davletova, S.; Rizhsky, L.; Liang, H.; Shengqiang, Z.; Oliver, D.J.; Coutu, J.; Shulaev, V.; Schlauch, K.; Mittler, R. Cytosolic Ascorbate Peroxidase 1 Is a Central Component of the Reactive Oxygen Gene Network of Arabidopsis. Plant Cell 2005, 17, 268–281, doi: 10.1105/tpc.104.026971.

17. Narendra, S.; Venkataramani, S.; Shen, G.; Wang, J.; Pasapula, V.; Lin, Y.; Kornyeyev, D.; Holaday, A.S.; Zhang, H. The Arabidopsis ascorbate peroxidase 3 is a peroxisomal membrane-bound antioxidant enzyme and is dispensable for Arabidopsis growth and development. Journal of Experimental Botany 2006, 57, 3033–3042, doi: 10.1093/jxb/erl060.

18. Bonekamp, N.A.; Völkl, A.; Fahimi, H.D.; Schrader, M. Reactive oxygen species and peroxisomes: struggling for balance. Biofactors 2009, 35, 346–355, doi: 10.1002/biof.48.

19. Costa, A.; Drago, I.; Behera, S.; Zottini, M.; Pizzo, P.; Schroeder, J.I.; Pozzan, T.; Schiavo, F.L. H2O2 in plant peroxisomes: an in vivo analysis uncovers a Ca2+-dependent scavenging system. The Plant Journal 2010, 62, 760–772, doi: 10.1111/j.1365-313X.2010.04190.x.

20. Foyer, C.H.; Noctor, G. Redox Homeostasis and Antioxidant Signaling: A Metabolic Interface between Stress Perception and Physiological Responses. The Plant Cell 2005, 17, 1866–1875, doi: 10.1105/tpc.105.033589.

21. Ahmad, P.; Sarwat, M.; Sharma, S. Reactive oxygen species, antioxidants and signaling in plants. Journal of Plant Biology 2008, 51, 167–173, doi: 10.1007/BF03030694.

22. Sies, H.; Berndt, C.; Jones, D.P. Oxidative Stress. Annu. Rev. Biochem. 2017, 86, 715–748, doi: 10.1146/annurev-biochem-061516-045037.

23. Mittler, R.; Vanderauwera, S.; Suzuki, N.; Miller, G.; Tognetti, V.B.; Vandepoele, K.; Gollery, M.; Shulaev, V.; Van Breusegem, F. ROS signaling: the new wave? Trends in Plant Science 2011, 16, 300–309, doi: 10.1016/j.tplants.2011.03.007.

24. Joo, J.H.; Bae, Y.S.; Lee, J.S. Role of Auxin-Induced Reactive Oxygen Species in Root Gravitropism. Plant Physiol. 2001, 126, 1055–1060, doi: 10.1104/pp.126.3.1055.

25. Shin, R.; Schachtman, D.P. Hydrogen peroxide mediates plant root cell response to nutrient deprivation. Proceedings of the National Academy of Sciences of the United States of America 2004, 101, 8827–8832, doi: 10.1073/pnas.0401707101.

26. Sarath, G.; Hou, G.; Baird, L.M.; Mitchell, R.B. ABA, ROS and NO are Key Players During Switchgrass Seed Germination. Plant Signal Behav 2007, 2, 492–493.

27. Liu, Y.; Ye, N.; Liu, R.; Chen, M.; Zhang, J. H2O2 mediates the regulation of ABA catabolism and GA biosynthesis in Arabidopsis seed dormancy and germination. Journal of Experimental Botany 2010, 61, 2979–2990, doi: 10.1093/jxb/erq125.

28. Potikha, T.S.; Collins, C.C.; Johnson, D.I.; Delmer, D.P.; Levine, A. The Involvement of Hydrogen Peroxide in the Differentiation of Secondary Walls in Cotton Fibers. Plant Physiol. 1999, 119, 849–858, doi: 10.1104/pp.119.3.849.

29. Li, S.-W.; Xue, L.; Xu, S.; Feng, H.; An, L. Hydrogen peroxide acts as a signal molecule in the adventitious root formation of mung bean seedlings. Environmental and Experimental Botany 2009, 65, 63–71, doi: 10.1016/j.envexpbot.2008.06.004.

30. Noriega, A.; Tocino, A.; Cervantes, E. Hydrogen peroxide treatment results in reduced curvature values in the Arabidopsis root apex. Journal of Plant Physiology 2009, 166, 554–558, doi: 10.1016/j.jplph.2008.07.009.

31. Cheng, H.; Liang, Q.; Chen, X.; Zhang, Y.; Qiao, F.; Guo, D. Hydrogen peroxide facilitates *Arabidopsis* seedling establishment by interacting with light signalling pathway in the dark. Plant Cell Environ 2019, 42, 1302–1317, doi: 10.1111/pce.13482.

32. Desikan, R.; A.-H.-Mackerness, S.; Hancock, J.T.; Neill, S.J. Regulation of the Arabidopsis Transcriptome by Oxidative Stress. Plant Physiology 2001, 127, 159–172, doi: 10.1104/pp.127.1.159.

33. Gadjev, I.; Vanderauwera, S.; Gechev, T.S.; Laloi, C.; Minkov, I.N.; Shulaev, V.; Apel, K.; Inze, D.; Mittler, R.; Van Breusegem, F. Transcriptomic Footprints Disclose Specificity of Reactive Oxygen Species Signaling in Arabidopsis. Plant Physiol. 2006, 141, 436–445, doi: 10.1104/pp.106.078717.

34. Mittler, R.; Vanderauwera, S.; Gollery, M.; Van Breusegem, F. Reactive oxygen gene network of plants. Trends in Plant Science 2004, 9, 490–498, doi: 10.1016/j.tplants.2004.08.009.

35. Hancock, J.; Desikan, R.; Harrison, J.; Bright, J.; Hooley, R.; Neill, S. Doing the unexpected: proteins involved in hydrogen peroxide perception. Journal of Experimental Botany 2006, 57, 1711–1718, doi: 10.1093/jxb/erj180.

36. Cheng, H.; Zhang, Q.; Guo, D. Genes that Respond to H2O2 Are Also Evoked Under Light in Arabidopsis. Mol. Plant 2013, 6, 226–228, doi: 10.1093/mp/sss108.

37. Wu, F.; Chi, Y.; Jiang, Z.; Xu, Y.; Xie, L.; Huang, F.; Wan, D.; Ni, J.; Yuan, F.; Wu, X.; et al. Hydrogen peroxide sensor HPCA1 is an LRR receptor kinase in Arabidopsis. Nature 2020, 578, 577–581, doi: 10.1038/s41586-020-2032-3.

38. Schubert, R.; Fischer, R.; Hain, R.; Schreier, P.H.; Bahnweg, G.; Ernst, D.; Heinrich, S.J. An ozone-responsive region of the grapevine resveratrol synthase promoter differs from the basal pathogen-responsive sequence. Plant Molecular Biology 1997, 34, 417–426, doi: 10.1023/A:1005830714852.

39. Guan, L.M.; Zhao, J.; Scandalios, J.G. Cis-elements and trans-factors that regulate expression of the maize Cat1 antioxidant gene in response to ABA and osmotic stress: H2O2 is the likely intermediary signaling molecule for the response. The Plant Journal 2000, 22, 87–95, doi: 10.1046/j.1365-313x.2000.00723.x.

40. Tsukamoto, S.; Morita, S.; Hirano, E.; Yokoi, H.; Masumura, T.; Tanaka, K. A Novel cis-Element That Is Responsive to Oxidative Stress Regulates Three Antioxidant Defense Genes in Rice. Plant Physiology 2005, 137, 317–327, doi: 10.1104/pp.104.045658.

41. Geisler, M.; Kleczkowski, L.A.; Karpinski, S. A universal algorithm for genome-wide in silicio identification of biologically significant gene promoter putative cis-regulatory - elements; identification of new elements for reactive oxygen species and sucrose signaling in Arabidopsis. The Plant Journal 2006, 45, 384–398, doi: 10.1111/j.1365-313X.2005.02634.x.

42. Veljovic-Jovanovic, S.; Noctor, G.; Foyer, C.H. Are leaf hydrogen peroxide concentrations commonly overestimated? The potential influence of artefactual interference by tissue phenolics and ascorbate. Plant Physiology and Biochemistry 2002, 40, 501–507, doi: 10.1016/S0981-9428(02)01417-1.

43. Cheeseman, J.M. Hydrogen peroxide concentrations in leaves under natural conditions. Journal of Experimental Botany 2006, 57, 2435–2444, doi: 10.1093/jxb/erl004.

44. Kwak, J.M.; Mori, I.C.; Pei, Z.-M.; Leonhardt, N.; Torres, M.A.; Dangl, J.L.; Bloom, R.E.; Bodde, S.; Jones, J.D.G.; Schroeder, J.I. NADPH oxidase AtrbohD and AtrbohF genes function in ROS-dependent ABA signaling in Arabidopsis. EMBO J 2003, 22, 2623–2633, doi: 10.1093/emboj/cdg277.

45. Lherminier, J.; Elmayan, T.; Fromentin, J.; Elaraqui, K.T.; Vesa, S.; Morel, J.; Verrier, J.-L.; Cailleteau, B.; Blein, J.-P.; Simon-Plas, F. NADPH oxidase-mediated reactive oxygen species production: subcellular localization and reassessment of its role in plant defense. Mol. Plant Microbe Interact 2009, 22, 868–881, doi: 10.1094/MPMI-22-7-0868.

46. Noctor, G.; Reichheld, J.-P.; Foyer, C.H. ROS-related redox regulation and signaling in plants. Semin. Cell Dev. Biol. 2017, doi: 10.1016/j.semcdb.2017.07.013.

47. Foyer, C.H.; Bloom, A.J.; Queval, G.; Noctor, G. Photorespiratory Metabolism: Genes, Mutants, Energetics, and Redox Signaling. Annu. Rev. Plant Biol. 2009, 60, 455–484, doi: 10.1146/annurev.arplant.043008.091948.

48. Kimura, M.; Yoshizumi, T.; Manabe, K.; Yamamoto, Y.Y.; Matsui, M. Arabidopsis transcriptional regulation by light stress via hydrogen peroxide-dependent and -independent pathways. Genes Cells 2001, 6, 607–617, doi: 10.1046/j.1365-2443.2001.00446.x.

49. Badger, M.R.; von Caemmerer, S.; Ruuska, S.; Nakano, H. Electron flow to oxygen in higher plants and algae: rates and control of direct photoreduction (Mehler reaction) and rubisco oxygenase. Philos Trans R Soc Lond B Biol Sci 2000, 355, 1433–1446.

50. Bailey, T.L.; Boden, M.; Buske, F.A.; Frith, M.; Grant, C.E.; Clementi, L.; Ren, J.; Li, W.W.; Noble, W.S. MEME SUITE: tools for motif discovery and searching. Nucleic Acids Research 2009, 37, W202–W208, doi: 10.1093/nar/gkp335.

51. Pavesi, G.; Zambelli, F.; Pesole, G. WeederH: an algorithm for finding conserved regulatory motifs and regions in homologous sequences. BMC Bioinformatics 2007, 8, 46–46, doi: 10.1186/1471-2105-8-46.

52. Kosugi, S.; Ohashi, Y. PCF1 and PCF2 Specifically Bind to Cis Elements in the Rice Proliferating Cell Nuclear Antigen Gene. Plant Cell 1997, 9, 1607–1619, doi: 10.1105/tpc.9.9.1607.

53. Trémousaygue, D.; Garnier, L.; Bardet, C.; Dabos, P.; Hervé, C.; Lescure, B. Internal telomeric repeats and ‘TCP domain’ protein-binding sites co-operate to regulate gene expression in Arabidopsis thaliana cycling cells. The Plant Journal 2003, 33, 957–966, doi: 10.1046/j.1365-313X.2003.01682.x.

54. Brandão, M.M.; Dantas, L.L.; Silva-Filho, M.C. AtPIN: Arabidopsis thaliana Protein Interaction Network. BMC Bioinformatics 2009, 10, 454, doi: 10.1186/1471-2105-10-454.

55. Demczuk, S.; Harbers, M.; Vennström, B. Identification and analysis of all components of a gel retardation assay by combination with immunoblotting. Proceedings of the National Academy of Sciences 1993, 90, 2574–2578.

56. Schultz, A.; Nottrott, S.; Watkins, N.J.; Lührmann, R. Protein-Protein and Protein-RNA Contacts both Contribute to the 15.5K-Mediated Assembly of the U4/U6 snRNP and the Box C/D snoRNPs. Mol Cell Biol 2006, 26, 5146–5154, doi: 10.1128/MCB.02374-05.

57. Dietz, K.-J.; Mittler, R.; Noctor, G. Recent Progress in Understanding the Role of Reactive Oxygen Species in Plant Cell Signaling. Plant Physiol 2016, 171, 1535–1539, doi: 10.1104/pp.16.00938.

58. Naydov, I.A.; Mudrik, V.A.; Ivanov, B.N. Light-induced hydrogen peroxide dynamics in protoplasts from leaves of both wild-type arabidopsis and its mutant deficient in ascorbate biosynthesis. Dokl Biochem Biophys 2010, 432, 137–140, doi: 10.1134/S1607672910030129.

59. Wen, F.; Xing, D.; Zhang, L. Hydrogen peroxide is involved in high blue light-induced chloroplast avoidance movements in Arabidopsis. Journal of Experimental Botany 2008, 59, 2891–2901, doi: 10.1093/jxb/ern147.

60. Mubarakshina, M.M.; Ivanov, B.N.; Naydov, I.A.; Hillier, W.; Badger, M.R.; Krieger-Liszkay, A. Production and diffusion of chloroplastic H2O2 and its implication to signalling. J. Exp. Bot. 2010, erq171, doi: 10.1093/jxb/erq171.

61. Beneloujaephajri, E.; Costa, A.; L’Haridon, F.; Métraux, J.-P.; Binda, M. Production of reactive oxygen species and wound-induced resistance in Arabidopsis thaliana against Botrytis cinereaare preceded and depend on a burst of calcium. BMC Plant Biology 2013, 13, 160, doi: 10.1186/1471-2229-13-160.

62. Brosché, M.; Overmyer, K.; Wrzaczek, M.; Kangasjärvi, J.; Kangasjärvi, S. Stress Signaling III: Reactive Oxygen Species (ROS). In Abiotic Stress Adaptation in Plants; Pareek, A., Sopory, S.K., Bohnert, H.J., Eds.; Springer Netherlands: Dordrecht, 2009; pp. 91–102 ISBN 978-90-481-3111-2.

63. Garretón, V.; Carpinelli, J.; Jordana, X.; Holuigue, L. The as-1 Promoter Element Is an Oxidative Stress-Responsive Element and Salicylic Acid Activates It via Oxidative Species. Plant Physiology 2002, 130, 1516–1526, doi: 10.1104/pp.009886.

64. Hudson, M.E.; Quail, P.H. Identification of Promoter Motifs Involved in the Network of Phytochrome A-Regulated Gene Expression by Combined Analysis of Genomic Sequence and Microarray Data. Plant Physiology 2003, 133, 1605–1616, doi: 10.1104/pp.103.030437.

65. Kosugi, S.; Suzuka, I.; Ohashi, Y. Two of three promoter elements identified in a rice gene for proliferating cell nuclear antigen are essential for meristematic tissue-specific expression. The Plant Journal 1995, 7, 877–886, doi: 10.1046/j.1365-313X.1995.07060877.x.

66. Welchen, E.; Gonzalez, D.H. Overrepresentation of Elements Recognized by TCP-Domain Transcription Factors in the Upstream Regions of Nuclear Genes Encoding Components of the Mitochondrial Oxidative Phosphorylation Machinery. Plant Physiol. 2006, 141, 540–545, doi: 10.1104/pp.105.075366.

67. Maruyama, K.; Todaka, D.; Mizoi, J.; Yoshida, T.; Kidokoro, S.; Matsukura, S.; Takasaki, H.; Sakurai, T.; Yamamoto, Y.Y.; Yoshiwara, K.; et al. Identification of Cis-Acting Promoter Elements in Cold- and Dehydration-Induced Transcriptional Pathways in Arabidopsis, Rice, and Soybean. DNA Res 2012, 19, 37–49, doi: 10.1093/dnares/dsr040.

68. Reichow, S.L.; Hamma, T.; Ferré-D’Amaré, A.R.; Varani, G. The Structure and Function of Small Nucleolar Ribonucleoproteins. Nucl. Acids Res. 2007, 35, 1452–1464, doi: 10.1093/nar/gkl1172.

69. Terns, M.P.; Terns, R.M. Small nucleolar RNAs: versatile trans-acting molecules of ancient evolutionary origin. Gene Expr. 2002, 10, 17–39.

70. Caparros-Ruiz, D.; Lahmy, S.; Piersanti, S.; Echeverría, M. Two Ribosomal DNA-Binding Factors Interact with a Cluster of Motifs on the 5′ External Transcribed Spacer, Upstream from the Primary Pre-rRNA Processing Site in a Higher Plant. European Journal of Biochemistry 1997, 247, 981–989, doi: 10.1111/j.1432-1033.1997.00981.x.

71. Sáez-Vasquez, J.; Caparros-Ruiz, D.; Barneche, F.; Echeverría, M. A Plant snoRNP Complex Containing snoRNAs, Fibrillarin, and Nucleolin-Like Proteins Is Competent for both rRNA Gene Binding and Pre-rRNA Processing In Vitro. Molecular and Cellular Biology 2004, 24, 7284–7297, doi: 10.1128/MCB.24.16.7284-7297.2004.

72. Zhou, H.; Chen, Y.-Q.; Du, Y.-P.; Qu, L.-H. The Schizosaccharomyces Pombe mgU6-47 Gene Is Required for 2′-O-Methylation of U6 snRNA at A41. Nucl. Acids Res. 2002, 30, 894–902, doi: 10.1093/nar/30.4.894.

73. Tycowski, K.T.; You, Z.-H.; Graham, P.J.; Steitz, J.A. Modification of U6 Spliceosomal RNA Is Guided by Other Small RNAs. Molecular Cell 1998, 2, 629–638, doi: 10.1016/S1097-2765(00)80161-6.

74. d’Orval, B.C.; Bortolin, M.-L.; Gaspin, C.; Bachellerie, J.-P. Box C/D RNA Guides for the Ribose Methylation of Archaeal tRNAs. The tRNATrp Intron Guides the Formation of Two Ribose-Methylated Nucleosides in the Mature tRNATrp. Nucl. Acids Res. 2001, 29, 4518–4529, doi: 10.1093/nar/29.22.4518.

75. Cavaillé, J.; Buiting, K.; Kiefmann, M.; Lalande, M.; Brannan, C.I.; Horsthemke, B.; Bachellerie, J.-P.; Brosius, J.; Hüttenhofer, A. Identification of Brain-Specific and Imprinted Small Nucleolar RNA Genes Exhibiting an Unusual Genomic Organization. PNAS 2000, 97, 14311–14316, doi: 10.1073/pnas.250426397.

76. Kishore, S.; Stamm, S. The snoRNA HBII-52 regulates alternative splicing of the serotonin receptor 2C. Science 2006, 311, 230–232.

77. Kishore, S.; Khanna, A.; Zhang, Z.; Hui, J.; Balwierz, P.J.; Stefan, M.; Beach, C.; Nicholls, R.D.; Zavolan, M.; Stamm, S. The snoRNA MBII-52 (SNORD 115) Is Processed into Smaller RNAs and Regulates Alternative Splicing. Hum. Mol. Genet. 2010, 19, 1153–1164, doi: 10.1093/hmg/ddp585.

78. Morlando, M.; Ballarino, M.; Greco, P.; Caffarelli, E.; Dichtl, B.; Bozzoni, I. Coupling between snoRNP assembly and 3|[prime]| processing controls box C/D snoRNA biosynthesis in yeast. The EMBO Journal 2004, 23, 2392–2401, doi: 10.1038/sj.emboj.7600254.

79. Miao, Y.; Jiang, L. Transient expression of fluorescent fusion proteins in protoplasts of suspension cultured cells. Nat. Protocols 2007, 2, 2348–2353, doi: 10.1038/nprot.2007.360.

80. Kiseleva, E.; Allen, T.D.; Rutherford, S.A.; Murray, S.; Morozova, K.; Gardiner, F.; Goldberg, M.W.; Drummond, S.P. A protocol for isolation and visualization of yeast nuclei by scanning electron microscopy (SEM). Nat. Protocols 2007, 2, 1943–1953, doi: 10.1038/nprot.2007.251.

81. Despres, C.; Subramaniam, R.; Matton, D.P.; Brisson, N. The Activation of the Potato PR-10a Gene Requires the Phosphorylation of the Nuclear Factor PBF-1. The Plant Cell 1995, 7, 589–598, doi: 10.1105/tpc.7.5.589.

82. Wang, X.; Bian, Y.; Cheng, K.; Zou, H.; Sun, S.S.-M.; He, J.-X. A Comprehensive Differential Proteomic Study of Nitrate Deprivation in Arabidopsis Reveals Complex Regulatory Networks of Plant Nitrogen Responses. J. Proteome Res. 2012, 11, 2301–2315, doi: 10.1021/pr2010764.

83. Ma, Y.; Yu, J.; Chan, H.L.Y.; Chen, Y.-C.; Wang, H.; Chen, Y.; Chan, C.-Y.; Go, M.Y.Y.; Tsai, S.-N.; Ngai, S.-M.; et al. Glucose-Regulated Protein 78 Is an Intracellular Antiviral Factor Against Hepatitis B Virus. Mol Cell Proteomics 2009, 8, 2582–2594, doi: 10.1074/mcp.M900180-MCP200.

84. Fujita, P.A.; Rhead, B.; Zweig, A.S.; Hinrichs, A.S.; Karolchik, D.; Cline, M.S.; Goldman, M.; Barber, G.P.; Clawson, H.; Coelho, A.; et al. The UCSC Genome Browser database: update 2011. Nucleic Acids Research 2010, 39, D876–D882, doi: 10.1093/nar/gkq963.

