## Supplemental Data for "The Identification of A ROS Responsive Motif that is regulated by snoRNP in *Arabidopsis*"

**Supplementary Table 1.** Primer pairs used for snoRNA detection.

| Z4 | CCTTGTTGATAAACTCTAATCTTC; CAGACATATGCTTGTCTATAATC |
| --- | --- |
| U59b | CTCTTATGATGTCAAAACTTTCG; ACTCTTCAGTTCAGGTCTCCA |
| R63 | GAGATATGATGATCAACAAATTGG; GAGAGATCAGAGTTTGGTTAAGAA |
| R88 | CTTCATCAGATAGTCGATGTGG; TGTAGCTGCTCAAGTTATGCTC |
| U25b | GGAAGAACAGTGATGAGTCAGTT; AGAAGAACTCAGTCCCTTAGATG |
| R74 | GCAGAGTTCTGTTGCAATTTG; CAATGTACCATCATGTACTGAATC |
| R38b | TGTGATGTTTTGGATGATGAG; GTGATGTGTGTCAGATTACAAAAG |
| contig2 | GAAAGTGATGATATGAAATTGTCG; GAATCAGACAAAAATAGTCAATCAC |
| contig5 | AAAGCATGGATCCGCCTAG; ACTCGGCTAAGAAGAAAGAAGA |
| contig7 | AGCCTGTGATGTTTGAGATTG; AAGCCTCAGAAAGTGGAGGT |
| R108 | TTATAGGGGAAATGAGGAATG; CAGATTGTATAAAAATTGGAGC |
| R86 | GAGCAAGGGCGTCTAGTTT; ACGAAAAACATATTTGATAGGC |
| Z43 | GGACAGTGACGATTGATATTAAG; AGGACTCAGAGTATGGGTGG |
| U24a | GGCCGGTGATGTAATCAA; GGCCTCAGAGATCTTGGTG |
| R41Y | CATGATGAGGATTGTTTTATTG; CACTCAGAATGAGTAGGAGGTA |
| R77 | GGCAACCTACTTGTTAATAAAAC; AATGTGCTTAAGACTACTAGACCA |
| R37 | GGATGTGGAAGGGGTTGA; TGGTGAAAGGGGAATTAATAC |
| R82 | GGCAGTGATGACTCGGAA; GGCTCAGATCAGCGGGGT |
| U49 | GACACAGAACGGTGGGAA; AACCCTCAGCGAACCCT |
| Z15 | TCAAGTGATGATTGATAACGC; CATCAGATAGGAGCGAAAGAC |
| AtU3 | CGACCTTACTTGAACAGGATCTGTTG; CTGTCAGACCGCCGTGCGT |
| AtU6 | GGACCATTTCTCGATTTATGCG; CAGGGAAGCCCCTGTAGGC |
| AtR82 | GCTTCTTTGATTGGGTC; GTGCCGGTAGATTAAGG |
| AtU14 | GCCGCCTAAGAGCTTTCGCC; TCAGACATCCAAGGAAGGATT |

**Supplementary Table 2.** The cloning and annotation of 18 identified snoRNAs. EST No. means clone amount in the sequenced library.

| Name | EST No. | family | Other name |
| --- | --- | --- | --- |
| snoR58y | 2 | C/D | R64 |
| Z4 | 3 | C/D | U27 |
| U59b | 3 | C/D |  |
| R63 | 3 | C/D |  |
| U25b | 3 | C/D |  |
| snoR38b | 2 | C/D | snoR38Y-1 |
| snoR108 | 1 | C/D |  |
| Z43 | 1 | C/D |  |
| snoR41Y | 1 | C/D | R7 |
| U24 | 1 | C/D | Z20 |
| snoR37-2 | 1 | C/D |  |
| U49 | 1 | C/D |  |
| R87 | 1 | C/D | Z107 |
| Z15 | 1 | C/D | snoR60 |
| snoR88 | 2 | H/ACA |  |
| snoR74 | 2 | H/ACA |  |
| snoR86 | 1 | H/ACA |  |
| snoR77 | 1 | H/ACA | Z7 |

**Supplementary Figure 1**

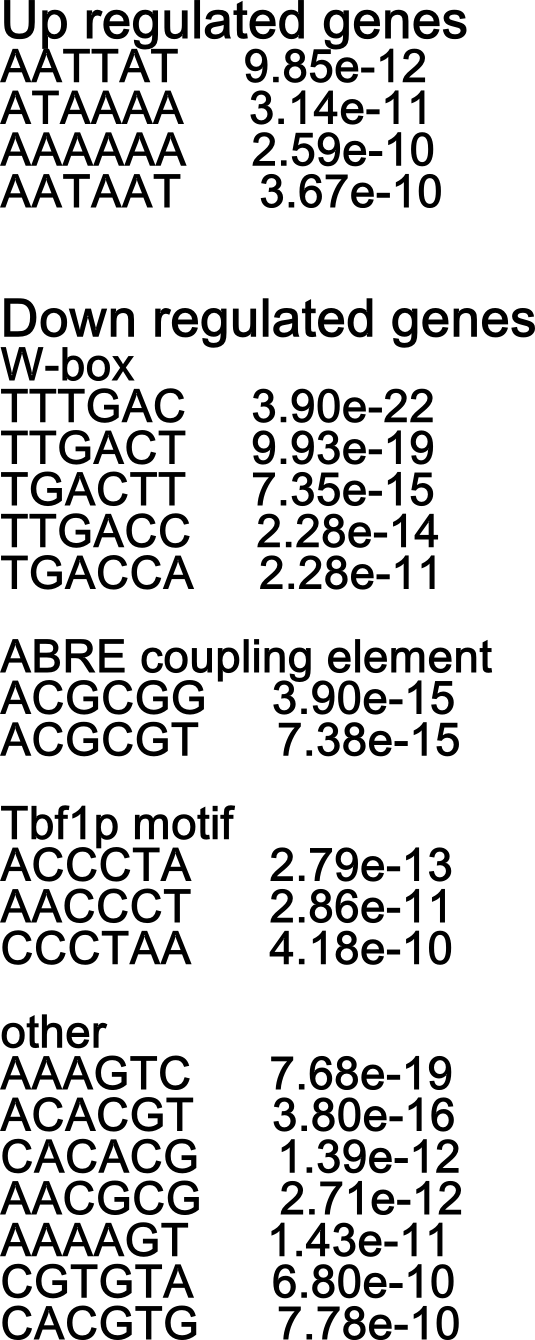

**Supplementary Figure 1**. The motifs identified using the microarray dataset GSE5530. Six .cel files (three H2O2 treated and three control) were analyzed using affy and limma package in R/Bioconductor (http://www.bioconductor.org). The -500 upstream sequences of H2O2 up-regulated and down-regulated genes were used for motif prediction using MEME Suite software against a background file consisted of words frequency in the 500 bp upstream sequences of all Arabidopsis genes' and TAIR Motif Analysis tool (http://www.arabidopsis.org/tools/bulk/motiffinder/index.jsp). No motif was predicted by the MEME software. This figure shows the results of TAIR Motif Analysis Tool.

**Supplementary Figure 2**

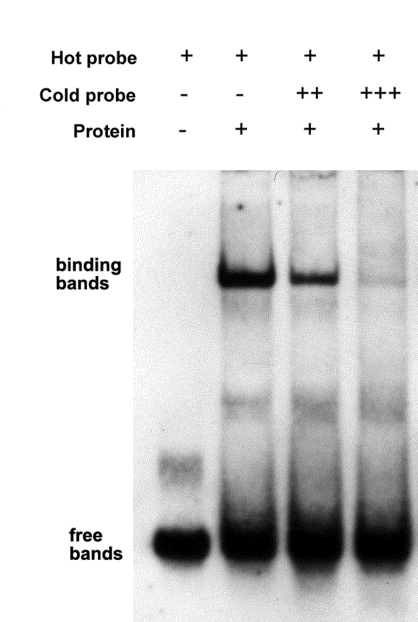

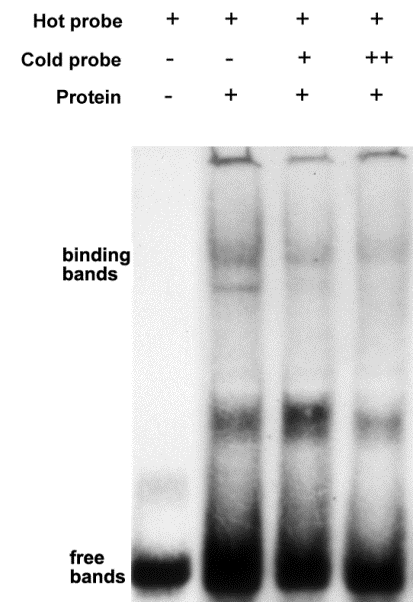

**A**

**B**

**Supplementary Figure 2.** Gel mobility shift assay for the validation of R-box in human (A) and mouse (B). Five microgram nuclear proteins from human HEK cells (A) or 20 microgram nuclear proteins from mouse H5V cells (B) were incubated with 1 ng 5'- Digoxigenin labeled R-box probe (lane 2-4, left to right) and then resolved by PAGE. Probes were transferred to positive charged nyon membrane and signals were detected with Digoxigenin detection kit. A negative control was performed by replacing nuclear protein with water (lane 1). A non-labeled cold probe was used as competitor in the binding reaction (1 ng lane 3 and 10 ng in lane 4). The signal bands of specific complexes and free non-binding probe were indicated as "binding bands" and "free bands" respectively.
